# Effects of mixing technique and ethanol removal on lipid nanoparticle physicochemical properties

**DOI:** 10.1101/2025.11.07.686408

**Authors:** Harsa Mitra, Titania Bethiana, Dongjie Jia, Mohammad Majidi, Pablo Mota-Santiago, Livia S. Manni, Izabela Milogrodzka, Kurt D. Ristroph, Arezoo M. Ardekani

## Abstract

Optimizing the production of lipid nanoparticle (LNP) therapeutics is necessary for drug delivery efficiency, stability, and scalability. A small but growing body of literature has begun to recognize that LNP properties (e.g., size, shape, and internal structure) depend on the flow conditions during mixing for antisolvent precipitation, in which LNPs are formulated. Here, we use different mixers, varying flow patterns (e.g., laminar or turbulent mixing) and flow rate ratios (FRR), i.e., 3:1 and 1:1, to prepare a standard LNP formulation. We then characterize the resulting formulations using small angle x-ray scattering (SAXS) to provide insights into particle shape/morphology, internal organization (L_*α*_ and H_*II*_ phases) of yeast RNA (yRNA), and structural differences/similarities that arise from the different mixing methods. The effect of ethanol removal on the LNPs’ structure, formulated from each mixing technique, is also discussed. We observed the 3:1 FRR mixers outperform the 1:1 configurations in certain desired LNP physiochemical properties. The differences observed in the LNPs produced across the two configurations are discussed. Furthermore, we use computational fluid dynamics to explain the turbulent mixing schemes among the 3:1 and 1:1 mixers.

## 1 INTRODUCTION

Development of nanocarrier systems for nucleic acid delivery, particularly mRNA-loaded lipid nanoparticles (LNPs), has opened new avenues for life-saving vaccines and precision genetic therapies. ^1–6^. These LNPs are self-assembled complexes that encapsulate an anionic nucleic acid payload through electrostatic complexation with cationic ionizable lipids, which co-precipitate with structural and stabilizing lipids ^1,7,8^. The complexation event occurs as part of a multistep assembly process that involves ethanol (solvent)– water (antisolvent) mixing, lipid supersaturation and precipitation, and steric stabilization by polyethylene glycol (PEG)-lipids on the surface of the resulting LNPs.

A critical, but often overlooked, factor influencing LNP structure and function is the nature of mixing during formulation. In early-stage laboratory settings (small scale), mixing is frequently carried out under laminar conditions using microfluidic techniques, or by hand pipetting ^9–11^.

In contrast, large-scale production relies on medium to large-scale turbulent mixing at higher flow rates to meet manufacturing demands ^12^. Recent studies have begun to show that these distinct hydrodynamic environments (laminar or turbulent or manual, and the specific geometry and operating conditions of each) significantly impact LNP performance, but a complete mechanistic understanding of how mixing affects LNP self-assembly remains unclear ^13–23^. Such an understanding is necessary when optimizing manufacturing processes, which in turn is critical to ensure efficient drug delivery/efficacy, formulation stability, and commercial scalability of the LNP modality ^13,16,23,24^.

Particle size, polydispersity, surface charge, RNBA encapsulation efficiency, and colloidal stability are typical critical quality attributes for LNPs. Additional physicochemical and structural attributes, including shape/sphericity and internal structural order, i.e., organization of the cargo and ionizable lipid, are emerging as key indicators of performance as well. ^16,23,25,25–34^. Recent studies using small-angle X-ray/neutron scattering (SAXS/SANS) have substantially probed LNP size, shape (surface to volume (S/V) ratios) and liquid crystalline (LC) phase behavior (order) and correlated these physiochemical properties to efficacy/in-vivo/vitro performance ^21,31,33,35–39^. The location and shape of Bragg peaks in SAXS coupled with cryo-electron microscopy have been utilized to emphasize the order of the LC phase. Interpretations of a shift from an isolated Bragg peak or lamellar, i.e., L*α*, feature to more complex diffraction peaks/patterns have been reported. These are evidence of a phase transition, such as a mixture/coexistence of phases, often from a L_*α*_ phase towards a non-lamellar, inverse hexagonal (H_*II*_) phase ^31,34,35^. Studies reported that this ordered internal structure of LNPs, formed by the nucleic acid cargo and ionizable lipid, tends to achieve improved therapeutic outcomes compared to those with more or less ordering ^19,31^. Separately, the presence of ‘bleb’-like structures, promoted by vesicle fusion, has also been associated with better transfection ^40,40,41^ . LC phase behavior in LNPs depends on material-driven factors such as selection of lipids, cargo, pH of the antisolvent, etc. ^26,31,40^ . Less attention has been paid to the mixing technique as a key contributor to the LNP internal organization/LC phase ^20,23,25,36^. Only one study to date has examined this directly, by comparing a coaxial turbulent jet mixer to a microfluidic method for producing LNPs. The authors reported that the coaxial mixer produced LNPs with smaller size, narrower size distributions, and higher encapsulation efficiency (EE) than LNPs made by microfluidics, but with a less ordered core ^20^. Another recent study reported that simply switching from hand mixing to microfluidics changed in vivo efficacy and organ tropism by orders of magnitude, but the internal structural ordering was not examined ^16^. Therefore a significant gap still remains in mechanistically connecting mixing regimes to LNP internal ordering, with all other compositional variables held constant.

The mixing regimes used in the literature can be broadly categorized as manual (pipetting), microfluidics, or turbulent. Microfluidic mixers such as the NanoAssemblr™ Ignite (Precision Nanosystems), which is a toroidal mixer operating in the laminar regime, have been instrumental in producing LNPs at lab scale. ^9,10,42^. Two major challenges in such mixers are, i) achieving efficient mixing, i.e., a more diffusion-dominant transport and mixing, at such low Reynolds numbers because of the low total flow rates (TFRs), and ii) scaling up for higher LNP production rates ^10,11^. Examples of turbulent mixers are the i) impinging jets mixers (IJM), such as the NOVA IJM (Helix, USA) and confined impinging jets (CIJ), and ii) multi-inlet vortex mixers (MIVM) ^23,43–46^ . The Pfizer/BioNTech SARS-CoV-2 vaccine is produced by impinging jets mixers, which facilitate rapid solvent exchange and flash nanoprecipitation for nanoparticle formulation. A major challenge with these mixers is operational cost, particularly at the lab scale, where the high volume of raw material required to perform a single run can be prohibitively expensive ^44^. For the CIJ, operational conditions need near-iso-momentum between opposing streams, constraining the flow rate ratio (FRR) to 1:1 and leading to a relatively high solvent fraction after mixing. High concentrations of ethanol are known to perturb lamellar spacing and affect internal order in LNPs ^47–49^. MIVMs have been reported to broaden residence-time distributions due to recirculation or backmixing when scaled up, causing wider size distributions (high polydispersity) ^50^.

Another critical process parameter in LNP production is the dialysis or diafiltration step used for solvent removal and/or buffer exchange following LNP formation to improve stability ^31,51–53^. This post-processing step can trigger fusion and structural rearrangements of LNPs (change of LC phase), especially as ionizable lipids shift protonation states during pH shifting ^40,41,51–53^.

In this pilot study, we use SAXS to investigate and compare the effects of the different mixing techniques and a dialysis step to remove ethanol while holding pH constant at 5. We compare hand mixing (HM), Ignite (laminar), CIJ, Helix, and MIVM (four different configurations). We quantify and discuss the observations from the multiple characterizations, in terms of LNP size, shape, internal structure (LC phase), and encapsulation efficiency. Computational fluid dynamics (CFD) is also utilized to visualize and quantify the turbulence mechanism and mixing across the two major high TFR mixers, CIJ and MIVM.

## 2 RESULTS AND DISCUSSION

### 2.1 Dynamic light scattering and encapsulation efficiency evaluation

DLS measurements, across Figs 1c and d, show that mixer selection significantly impacts LNP size, PDI, and EE for a given combination of lipids and RNA. These are quantified in Table 1. In terms of size, except for the hand-mixed, the rest of the 3:1 FRR mixers produced smaller diameter particles compared to the 1:1 FRR mixers. This trend was observed across both the non-dialyzed and dialyzed LNPs. Also, it can be observed in Figs 1c, except for hand mixed, CIJ, MIVM 2:2 adjacent and opposite, all other dialyzed LNPs had an increase in particle size compared to their non-dialyzed forms. The largest growth in LNP size post dialyzing was observed for the Helix mixer (12.43% increase). Across dialyzed/non-dialyzed, Ignite produced the smallest LNPs (62.53 nm), followed by the two MIVM 3:1 configurations. Among the dialyzed LNPs, MIVM 2:2 (adjacent) produced the largest particle size (105.5 nm), followed by the CIJ and MIVM 2:2 (opposite). For the final dialyzed particles, CIJ had the largest size of 97.38 nm.

**TABLE 1.**
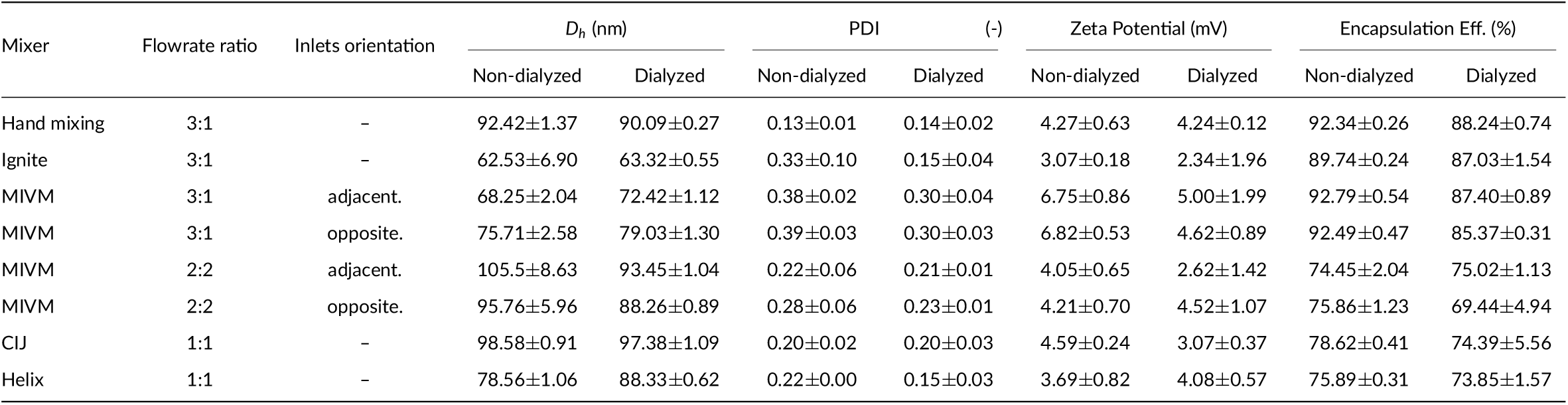
Dynamic light scattering (DLS) and RiboGreen assay results for LNPs produced with different mixers. *D*_*h*_ is the hydrodynamic diameter of the LNPs.

**FIGURE 1.**
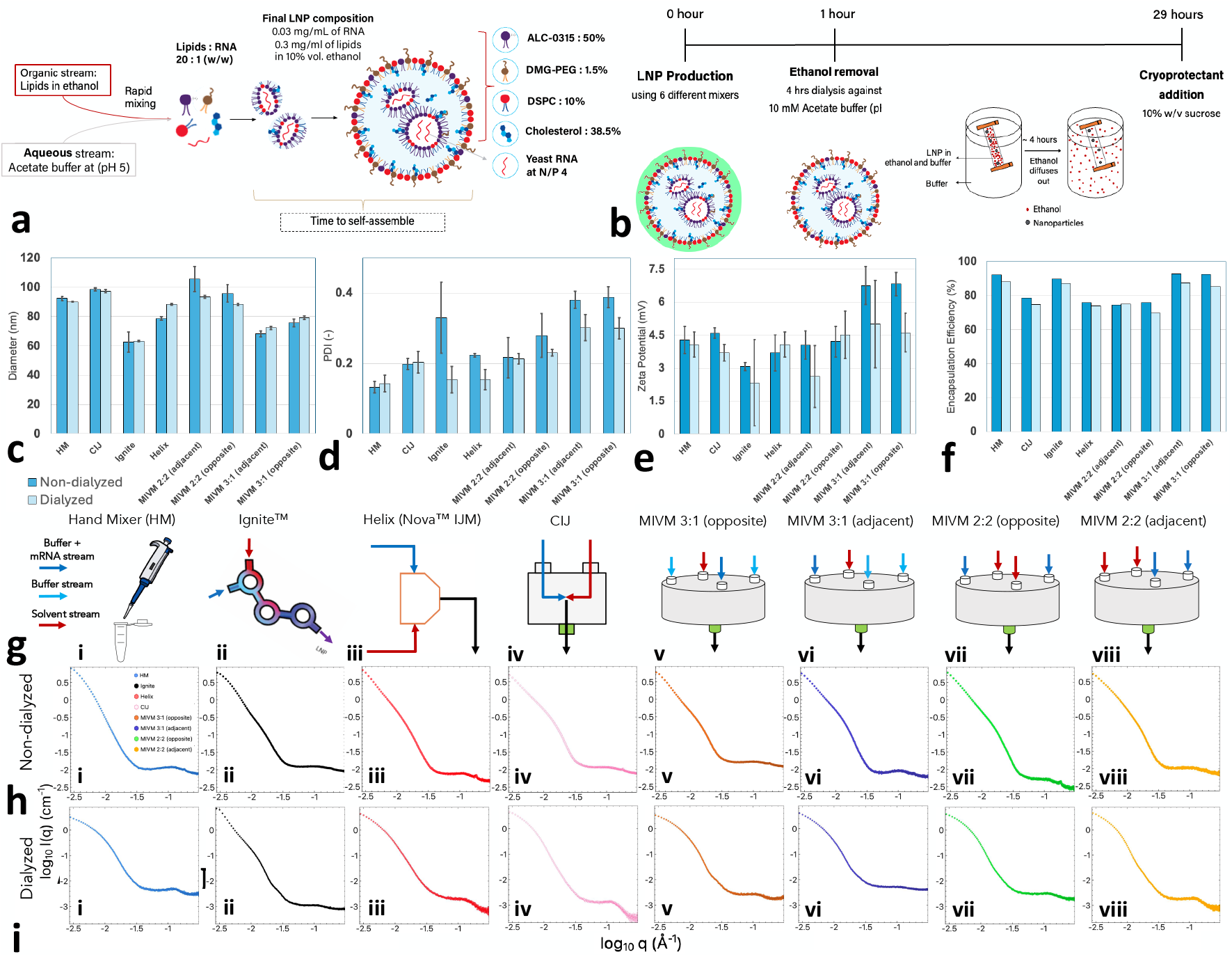
Self-assembly process, preparation methodology, and DLS and SAXS evaluations of the LNP formulations: a) Schematic of the selfassembly process utilized for producing the LNPs in this study. b) The process involved for producing the final LNP samples cryo-preserved with 10% w/v sucrose, after 29 hrs since LNPs are collected from each mixing technique. DLS-based, c) LNP size, d) polydispersity (PDI) index, and e) zeta potential. f) Encapsulation efficiency of the LNPs based on the ribogreen assays. g) The various different mixing strategies/techniques used. SAXS profiles for all tested LNPs from eighth different mixing techniques h) pre-dialysis and i) post-dialysis. Schematics are made using a combination of Microsoft Powerpoint and Biorender.com.

In terms of polydispersity (See Fig. 1d and Table 1), both the MIVM 3:1 configurations produced LNPs with the highest PDI values. Ignite had a similar performance with a PDI of 0.33, but a significant drop in post-dialysis polydispersity to 0.15. This decrease in PDI was non-monotonic across all the mixers, though a significant decrease in PDI was observed for Helix, MIVM 2:2 (opposite), and both the MIVM 3:1 configurations. The hand-mixed LNPs had the lowest PDI across all the cases. All the LNPs were neutral as the zeta potentials were well within +10 mV, as observed in Fig. 1e.

Finally, the encapsulation efficiency (EE), post mixing, the hand mixed, Ignite, and both the MIVM 3:1 configurations had EE greater than 89%. The hand-mixed technique had the highest EE across both pre (92.34%) and post dialyzing (89.74%). On the other hand, CIJ, Helix, and MIVM 2:2 configurations had lower than 80% EE (See Fig. 1d and Table 1). We observed a similar trend for all the LNPs post dialyzing, with the better performers, i.e., HM, Ignite, both MIVM 3:1 configurations, showing EE higher than 85%. This noticeable reduction in EE% can be observed for all the cases, except for MIVM 2:2 (adjacent) configuration.

### 2.2 SAXS evaluation

One-dimensional, *I*(*q*), curves from the SAXS measurements were reduced and their backgrounds subtracted (See Sec. 4.6 for details), for the non-dialyzed and dialyzed LNPs, from eight different mixing techniques, as shown in Figs. 1d and e, respectively. The Porod region is defined in the q-range, after the Guinier region (q > 0.01 Å^−1^ and) before the liquid crystalline (LC) phase (q < 0.1 Å^−1^). The scattering here decays as *I*(*q*) ∝ *q*^−*p*^, where *p* is the Porod exponent that is dependent on particle shape ^54–57^. For the non-dialyzed LNPs, we observe *p* to be within 3.03 to 3.31 for Ignite, Helix, CIJ, and the MIVM 2:2 configurations, indicating rough, heterogeneous interfaces ^56^. LNPs produced by hand mixing exhibited a *p* = 4 and LNPs made by MIVM 3:1 configurations had 3.4, suggesting slightly smoother boundaries at the LNP/buffer interface, especially for the hand mixed LNPs. Once dialyzed, all the mixing configurations had a *p* > 3.8. This is consistent with compact particle shapes possessing sharper, smoother LNP interfaces.

To better visualize the liquid crystalline (LC) phase, *I*(*q*) for the *q*, where *q* is the reciprocal space, i.e., *q* = 2*π*/*d*_0_, range around the primary Bragg peak (*n* = 1) is presented across Figs. 2a and b, for the non-dialyzed and dialyzed cases. The legends for each mixer across Figs. 1h,i, and Figs. 2a,b, are the same. The LC phase has been reported to arise around a *q* of 0.1 Å^−1^, due to the mRNA and ionizable lipid complexation and arrangement within the LNPs ^8,31,35,53,58^. Based on the existence of single or multiple Bragg peaks at or after 0.1 Å^−1^, reports on the phase of the order/packing of the LC phase have been studied ^31,35,39^. Although in the case of the non-dialyzed LNPs, we observed the existence of two major peaks, with the secondary peak (*n* = 2) forming after the primary LC phase peak (See Fig. 2a). This peak disappears post-dialysis as observed in Fig. 2b. It is also interesting to note, the sharpness of the *n* = 2 Bragg peak in Figs. 2a as compared to the primary peaks (*n* = 1). Interestingly, *n* = 2 occurs at approximately 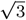times *n* = 1 peak location. This is exemplary of 2nd-order reflection in an inverse hexagonal, H_*II*_, phase ^39^. Such observation concludes that the non-dialyzed LNPs have stronger internal Bragg features (possibly both L_*α*_/H_*II*_), but after dialysis, once the ethanol is removed, the secondary reflections disappear, suggesting a reduction in the order of the (L_*α*_/H_*II*_) phase ^31^.

**FIGURE 2.**
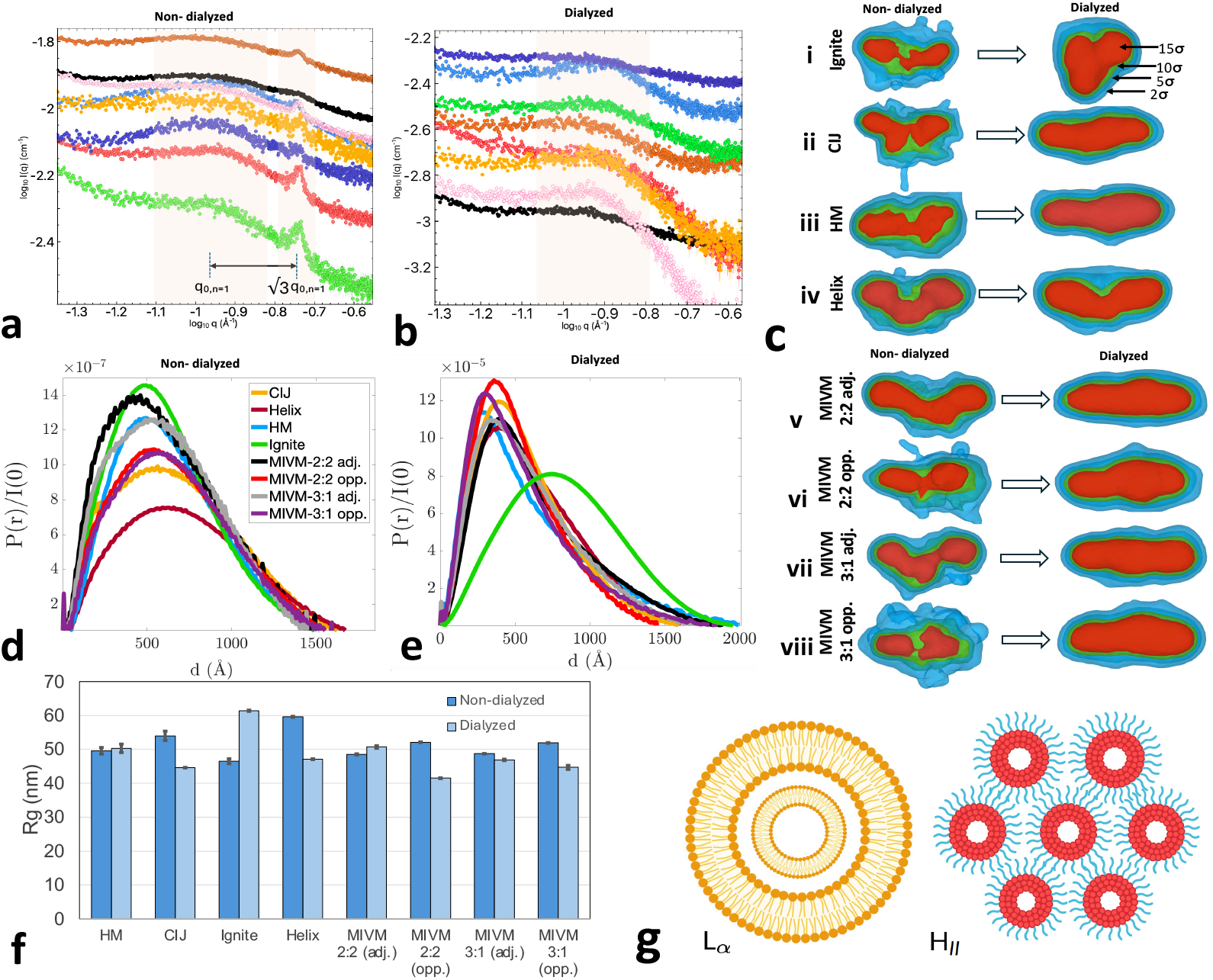
Liquid crystalline (LC) phase of the LNPs, a) non-dialyzed and b) dialyzed. Shaded regions represent the broad profile of the primary (*n* = 1) and secondary (*n* = 2) Bragg peaks. The Bragg peak ratio, i.e., 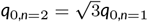 is overlaid in part a. The legends for each mixer here are the same as in Figs. 1d and e. c) DENSS reconstruction of both the non-dialyzed and dialyzed LNPs from each mixing approach. The color contours represent 15σ (red), 10σ (green), 5σ (cyan), and 2σ (blue). PDDF (GNOM) from the different mixers for d) non-dialyzed and e) dialyzed. f) Radius of gyration (*Rg*) for each mixing case for both non-dialyzed and dialyzed LNPs from the GNOM fitting. g) Full section of LNPs representing a fully lamellar (left) and inverse hexagonal phase (right). Schematics are made using a combination of Microsoft PowerPoint and Biorender.com.

In Figs. 2a and b, slight shifts and the broadness/narrowness of the primary peaks can be observed across the *I*(*q*) curves among the LNPs from the different mixers as well as pre- and post-dialyzing. These Bragg peaks arising due to the (L_*α*_/H_*II*_) phase, i.e., internal order of the LNPs, are quantified using a shape-independent model, i.e., Broad-Lorentzian (BL) model (See Eq. 1). In order to quantify the degree of order of the LC phase, the intensity, width, and area under the curve (AUC) of the mRNA/ionizable lipid Bragg peak have also been reported ^21,31^. The quantified parameters are presented for the dialyzed and non-dialyzed cases in Tables 2 and 4. Additionally, just the location of the secondary peak observed for the non-dialyzed LNPs is also reported in Table 3.

**TABLE 2.** Summary of fitted parameters for non-dialyzed LNPs from the different mixing techniques.

| Type | $\chi^2$ (-) | Peak Location ($2\pi/q_0$) [Å] | $m$ (-) | $\xi$ [Å] | $n$ (-) |
| --- | --- | --- | --- | --- | --- |
| Ignite | 1.05 | 64.77 | 2.65 | 18.65 | 1.32 |
| Hand mixing | 1.19 | 55.75 | 3.02 | 26.02 | 1.02 |
| CIJ | 1.01 | 57.64 | 1.94 | 20.16 | 0.62 |
| Helix | 0.99 | 57.64 | 1.66 | 11.85 | 2.27 |
| MIVM 2:2 (opposite) | 0.82 | 57.20 | 1.03 | 20.17 | 0.84 |
| MIVM 2:2 (adjacent) | 0.88 | 57.15 | 0.65 | 19.71 | 0.73 |
| MIVM 3:1 (opposite) | 0.99 | 61.59 | 2.47 | 21.94 | 0.18 |
| MIVM 3:1 (adjacent) | 0.98 | 61.01 | 2.67 | 26.63 | 0.82 |

**TABLE 3.** Summary of fitted parameters for non-dialyzed secondary peak of LNPs from the different mixing techniques.

| Type | $\chi^2$ (-) | Peak Location ($2\pi/q_0$) [Å] |
| --- | --- | --- |
| Ignite | 2.22 | 43.63 |
| Hand mixing | 3.04 | 37.85 |
| CIJ | 2.64 | 38.07 |
| Helix | 2.45 | 37.17 |
| MIVM 2:2 (opposite) | 1.53 | 37.85 |
| MIVM 2:2 (adjacent) | 0.90 | 34.33 |
| MIVM 3:1 (opposite) | 1.21 | 37.17 |
| MIVM 3:1 (adjacent) | 1.26 | 40.02 |

**TABLE 4.** Summary of fitted parameters for dialyzed LNPs from the different mixing techniques.

| Type | $\chi^2$ (-) | Peak Location ($2\pi/q_0$) [Å] | $m$ (-) | $\xi$ [Å] | $n$ (-) |
| --- | --- | --- | --- | --- | --- |
| Ignite | 1.07 | 61.00 | 2.48 | 38.05 | 0.74 |
| Hand mixing | 0.91 | 59.55 | 3.31 | 19.03 | 0.04 |
| CIJ | 0.90 | 60.41 | 2.49 | 15.05 | 0.02 |
| Helix | 1.07 | 62.45 | 2.35 | 11.27 | 3.60 |
| MIVM 2:2 (opposite) | 1.04 | 53.24 | 1.77 | 16.88 | 0.92 |
| MIVM 2:2 (adjacent) | 0.84 | 62.14 | 3.02 | 17.04 | 0.64 |
| MIVM 3:1 (opposite) | 0.99 | 62.27 | 3.03 | 21.2 | 2.11 |
| MIVM 3:1 (adjacent) | 1.04 | 59.83 | 2.79 | 23.01 | 2.60 |

As quantified by the BL model, for the non-dialyzed and dialyzed LNPs, the primary peak (*n* = 1) location is found to be around 55.75 - 64.77 Åand 53.24 - 62.45 Å, respectively. The secondary peak (*n* = 2) varied with a range of 37.17 - 43.63 Å. This shows a 16.18% (non-dialyzed) and 17.30% (dialyzed) variation for the internal organization of the yRNA and ALC-0315 across the various mixing configurations. Furthermore, we utilize the BL model parameters to elucidate the details of the internal order of the LC phase in the LNPs. For non-dialyzed LNPs, hand mixing (HM) produces the most tightly packed LC phase (a mixture of L_*α*_/H_*II*_) with a spacing of 55.75 Å. This can be additionally quantified using the Lorentz exponent, *m*, in the BL model (See Eq. 1), which describes the shape and sharpness of the Bragg peak. For HM LNPs, *m* is found to be 3.02 (See Table 2), indicating the most well-defined short-range order among all the non-dialyzed LNP cases. This possibly suggests that slower convective timescales (See Sec. 2.3) could be favorable for better core packing, though the flow regime in the hand mixing technique, i.e., laminar/turbulent or a mixture of both, is difficult to characterize. Additionally, the correlation length (*ξ*), within the LC phase context, describes the extension of these ordering coherent domains within the LNPs. For the HM technique, the extension of this LC domain is found to be for *ξ* = 26.02 Å. Ignite and both the MIVM 3:1 configurations showed *m* values more than 2.4, suggesting less-defined LC phase ordering. Among the 1:1 FRR mixers, CIJ showed the best short-range order with a *m* of 1.94, followed by Helix (which had the lowest LC domain across all the mixers). The MIVM 2:2 configurations had the lowest *m* values, suggesting a disordered LC phase among all the mixers.

Post dialyzing, the LC phase (lack of H_*II*_) ordering trend followed (See Table 4), HM>MIVM 3:1 (opposite)>MIVM 2:2 (adjacent)> MIVM 3:1 (adjacent)>CIJ>Ignite>MIVM 2:2 (opposite). It is also important to note the reduction in the significant difference observed across the *m* values, for the 3:1 and the 1:1 FRR configurations of the MIVM across the non-dialyzed LNPs. Especially among the MIVM 2:2 (adjacent) and MIVM 3:1 (opposite). This observation suggests re-ordering and the importance of the dialysis step for the order of the LC domain.

Shape analysis was performed on all the LNPs and both the non-dialyzed and dialyzed cases as shown in Fig. 2c-f. We utilize the algorithm DENSS (DENsity from Solution Scattering), algorithm can be applied to 1d-SAXS data (*I*(*q*)) from the LNPs to reconstruct ab initio electron density in three dimensions ^26,59,60^. See Sec. 4.6 for details on the DENSS methodology. The 3D-volume reconstructions are presented in Figs. 2c. We observe a core-shell structure with the electron-rich yRNA within the core of the particles, surrounded by the lipids, which have lower contrast. Post-dialyzing, we observe that all eight LNPs from each mixer had an ellipsoidal shape except for the Ignite (laminar with 3:1 FRR), which was comparatively more spherical. As compared to the non-dialyzed LNPs, we notice the shape and the yRNA/lipid regions gaining more homogeneity after dialysis is performed (See Figs. 2c). This supports the Porod exponents of *p* < 3.4 for the non-dialyzed cases, except for the hand mixed, which showed smooth interfaces with a *p* = 4. Additionally, it also points to the importance of the dialysis step. In this study, we did not observe any “bleb”-like structures.

The pairwise distance distribution function (PDDF) was calculated using GNOM (BioXTAS RAW) utilized to get the *P*(*r*) functions (See Figs. 2d and e) and quantify the radius of gyration (*Rg* ) for each LNP dispersion. Details about the fitting techniques can be found in Sec. 4.6. The quantified *Rg* for each LNP is presented in Fig. 2f. We observed a significant increase in *Rg* only in the cases of Ignite post dialysis (See Fig. 2f). A considerable reduction in size was observed for all other cases, except for hand mixing, and both the MIVM adjacent configurations. These results are tabulated in Table. 5. Similar observations have been reported previously for microfluidic mixing, where an increased size after dialysis, particularly when nucleic acid cargo was present ^52,53^. In contrast, turbulent mixers (e.g., MIVM) generate particles that are closer to equilibrium upon formation, and hence, the little or no size increase post-dialysis ^31^. Notably, these structural changes were not reflected in the hydrodynamic diameters measured by DLS, which remained constant or slightly decreased (increases only for Helix) (See Fig. 1c).

The PDDF confirmed the existence of the spherical LNP shape (See Fig. 2e) for the Ignite case, which was previously observed through the DENSS reconstruction. Except for CIJ, Helix, MIVM 2:2 (opposite), an increase in *Dmax* was observed for the LNPs post dialysis (See Table 5). Before dialyzing, Helix-based LNP showed the highest *Dmax* of 195 nm and *Rg* = 59.6 nm. Post-dialysis, the largest *Dmax* was observed for the hand-mixed LNPs with 205 nm (*Rg* = 50.3 nm). It is important to note, all the LNPs characterized through the PDDF estimate larger sizes as compared to the quantified diameters (*D*_*h*_), i.e., sphere-equivalent hydrodynamic diameter, from DLS measurements (See Table. 1). But it is also important to consider non-spherical shapes, as in our case, direct size (diameter) comparisons can be misleading if proper *R*_*g*_ to *R*_*h*_ conversions are not performed ^61^.

**TABLE 5.** Results of the inversion performed using GNOM for extracting the PDDF.

| Type | $R_g$ (nm) | Error (nm) | $D_{max}$ (nm) | Total estimate (-) | $\chi^2$ (-) | $I(0)$ (cm<sup>-1</sup>) |
| --- | --- | --- | --- | --- | --- | --- |
| HM-ND | 49.6 | 0.96 | 160 | 0.69 | 1.22 | 81.11 |
| HM-D | 50.3 | 1.25 | 205 | 0.66 | 0.95 | 55.50 |
| CIJ-ND | 54.0 | 1.38 | 170 | 0.67 | 1.10 | 82.30 |
| CIJ-D | 44.6 | 0.09 | 160 | 0.78 | 6.75 | 62.43 |
| Ignite-ND | 46.5 | 0.73 | 155 | 0.61 | 1.54 | 72.68 |
| Ignite-D | 61.4 | 0.17 | 180 | 0.66 | 7.07 | 18.35 |
| Helix-ND | 59.6 | 2.06 | 195 | 0.67 | 1.12 | 19.20 |
| Helix-D | 47.1 | 0.12 | 160 | 0.70 | 1.08 | 66.70 |
| MIVM (2:2 adj)-ND | 48.5 | 0.18 | 150 | 0.60 | 0.94 | 68.18 |
| MIVM (2:2 adj)-D | 50.7 | 0.39 | 190 | 0.68 | 0.77 | 69.59 |
| MIVM (2:2 opp)-ND | 52.1 | 0.10 | 160 | 0.72 | 1.62 | 82.60 |
| MIVM (2:2 opp)-D | 41.5 | 0.12 | 145 | 0.72 | 1.13 | 55.20 |
| MIVM (3:1 adj)-ND | 48.7 | 0.09 | 145 | 0.58 | 1.69 | 73.34 |
| MIVM (3:1 adj)-D | 46.9 | 0.21 | 170 | 0.65 | 1.00 | 62.04 |
| MIVM (3:1 opp)-ND | 51.9 | 0.10 | 160 | 0.63 | 1.62 | 85.23 |
| MIVM (3:1 opp)-D | 44.7 | 0.60 | 190 | 0.63 | 0.97 | 52.43 |

Furthermore, we utilize the Volatility of Ratio (Vr) (See Eq. 2 in Sec. 4.6) method, which describes the similarity/variability in the ratio of scattering intensity, i.e., *I*(*q*), between two different LNP systems ^62^.

Vr has been reported to be more robust than other methods of profile comparison for SAXS profiles. The Vr method was applied to all the LNP cases, and the resulting similarities are presented as a heat map in Figs. 3a-d. We perform separate similarity analysis for the LC phase for the q range of 0.08 - 0.2 Å^−1^ (See Figs. 3c and d). For the non-dialyzed LNPs in Fig. 3a, based on the whole q-range, we observe the MIVM 2:2 (opposite) to be dissimilar to all the other mixers, except for the Helix, which shows good similarity to the MIVM 2:2 (opposite). The other strong similarity can be noticed between the MIVM 3:1 (opposite), Ignite, and the CIJ. Interestingly, when the q-range around the LC phase is considered (See Fig. 3d), we still observe strong similarity between MIVM 3:1 (opposite), Ignite, and CIJ. In Fig. 3c, both the configurations of MIVM 2:2 and Helix had the most dissimilar SAXS profile as compared to the rest.

**FIGURE 3.**
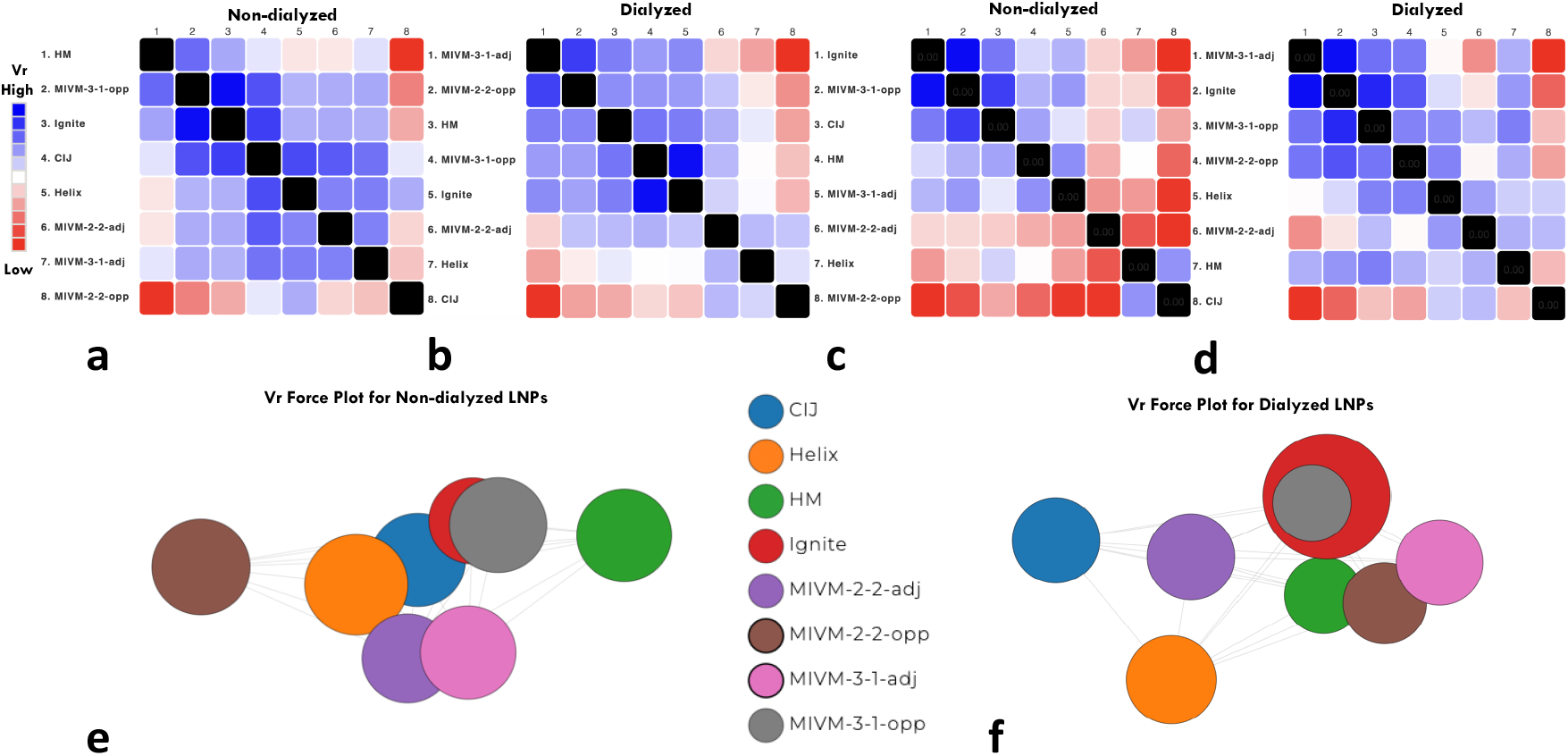
Volatility of Ratio (Vr) ^62^ of SAXS curves from all mixers. Vr calculated for the whole q-range a) non-dialyzed and b) dialyzed. Vr for the q-range of 0.08 to 0.2 Å^−1^, c) non-dialyzed and d) dialyzed. Vr color ranges for a) 8.22 (blue)-1.60 (red), b) 6.95 (blue)-0.94 (red), c) 1.49 (blue)-0.25 (red), and d) 3.66 (blue)-0.66 (red). Clustered force plots of all the mixers for e) non-dialyzed and f) dialyzed.

Once the LNPs are dialyzed (See Fig. 3b), we observe strong trends between Ignite, MIVM 3:1 (opposite), and MIVM 2:2 (opposite). The CIJ was dissimilar to all the mixers, except for some weak similarity to Helix and MIVM 2:2 (adjacent). We noticed the strongest dissimilarity between the CIJ and the MIVM 3:1 (adjacent). In Fig. 3d, LC phase qregion, other than the Helix and MIVM 2:2 (adjacent), CIJ performed the most dissimilar to the rest. It should be noted, once again, the MIVM 3:1 (adjacent) and CIJ were the most dissimilar pair of mixers, which suggests strong differences in the LC phase behavior. Also, similar to the whole SAXS q-range behavior, the LC phase SAXS profiles are also very similar between the two MIVM 3:1 (opposite/adjacent) and Ignite. This suggests the existence of strong differences between the 3:1 and 1:1 FRR mixers. Especially between the CIJ/Helix and the MIVM 3:1 configurations.

This divergence between the 1:1 and 3:1 FRR mixers (both laminar and turbulent) could be attributed to both the changes in solvent quality and the shear/flow profile differences across the mixers. Especially, between CIJ/Helix and the MIVMs, strongly different turbulent flows occur at the operating conditions in this study. We will talk in more detail on the fluid dynamic aspects in Sec. 2.3. Additionally, the solvent quality effects can be observed between the MIVMs, i.e., 3:1 and 2:2. But it can also be noticed that the MIVM 2:2 configurations and Helix outperform the CIJ, in being similar to the 3:1 FRR mixers, Ignite, HM, post dialyzing (See Fig. 3b). In addition to the heat maps, force plots for each LNPs based on their SAXS profiles have been presented in Figs. 3e and f. Closely clustered circles represent highly similar LNPs (based on their Vr from SAXS curves) and larger circles signify the magnitude of their *Rg* . Before dialyzing, as seen in Fig. 3e, most formulations (CIJ, Helix, Ignite) formed a similarity cluster at the center, suggesting comparable structures, while the hand-mixed and certain MIVM 2:2 (adjacent) deviate. Dialyzing appears to amplify the difference among the helix, CIJ, and MIVM 2:2 (adjacent), as compared to the Ignite, hand mixed, MIVM 3:1 configurations, and MIVM 2:2 (opposite) (See Fig. 3f).

### 2.3 Computational fluid dynamics evaluation

In this section, we discuss the high-fidelity CFD results to demonstrate the differences in turbulence and mixing of two very contrasting mixers, i.e., the CIJ and MIVM. As discussed before in Sec. 2.2, here we will explain the differences observed in the LNPs produced across the CIJ and MIVM from the fluid flow dynamics viewpoint. To complement the experimental results, we simulated two flow cases: the CIJ with a 1:1 flow rate ratio with a TFR of 120 mL/min and the MIVM 2:2 (opposite), i.e., with opposing streams, and a TFR of 80 mL/min. Both the 1:1 FRR configurations were selected to keep the solvent conditions the same across the two mixers. The results from the CFD simulations are summarized in Figure 4. Details on the CFD technique are provided in Sec. 4.7. The turbulent kinetic energy, *k*, measures the intensity of the turbulence. The mixing index, *MI*, measures the homogeneity of mixing across the horizontal plane. The vertical position is non-dimensionalized using *Z*^∗^ = *Z*/*D*, where *D* is the cross-section diameter. The size of the mixers in Figure 4 is scaled to the *Z*^∗^ axis of each plot.

**FIGURE 4.**
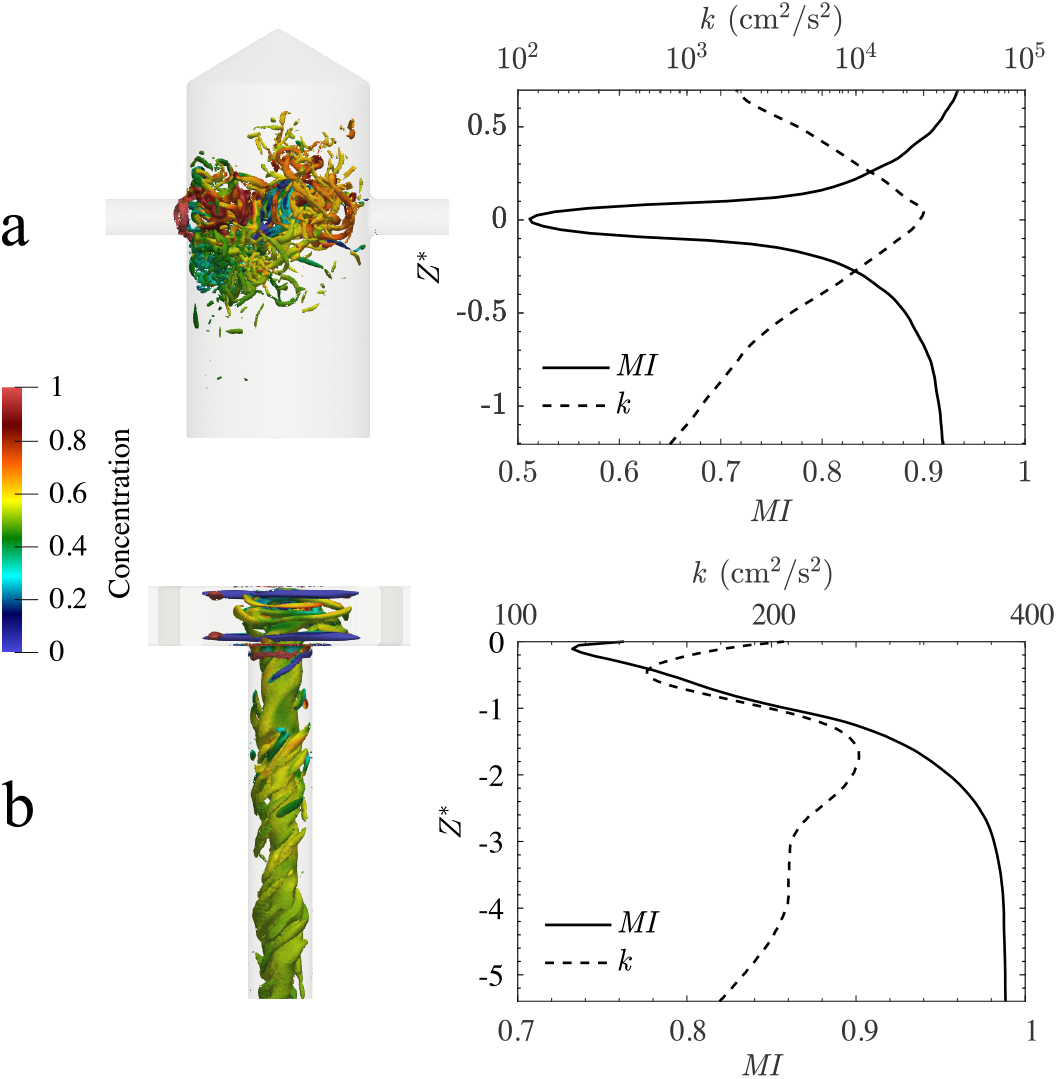
Q-criterion (left) contour colored with the solvent, i.e., ethanol, concentration and change of mixing index (right), *MI*, (solid line) and average turbulent kinetic energy, *k*, (dashed line) along the dimensionless vertical position *Z*^∗^ for (a) the CIJ and b) the MIVM 2:2 (opposite). It should be noted that the CIJ generates two orders of magnitude higher turbulent kinetic energy, as compared to the MIVM under the same 1:1 FRRs.

The turbulence in the MIVM 2:2 (opposite) outlet channel forms a turbulent tube with swirling turbulent structures. The mixing is near complete at the onset of the turbulence, as shown in the *Q*–criterion results colored with the solvent (ethanol) concentration (See Figs. 4a and b). Here Q-criterion should not be confused with “q” which defines the reciprocal space in the SAXS results. The turbulence persists downstream of the outlet tube with a small decay in energy. On the other hand, the CIJ produces strong turbulence at the flow impinging point. This results in the turbulent kinetic energy at its highest at the location where the fluids are least mixed (a hundred times more than the maximum *k* quantified in the MIVM). This is demonstrated in Figure 4b, where at the plane of impingement (*Z*^∗^ = 0), the mixing index is at the minima while the turbulent kinetic energy is at the maxima. The turbulence structure in the CIJ is more chaotic both spatially and temporally, with no consistent structure. Furthermore, the turbulence energy exhibits a strong power-law decay towards the downstream. Despite similar operating conditions and industrial applications, the turbulence and mixing dynamics of MIVM and CIJ are significantly different, as demonstrated in the CFD results. These fluid dynamical differences can provide foundations for theories on the variation in production performance presented in Sec. 2.2 (For more detailed discussion see ^63^).

## 3 CONCLUSIONS

In this study, we demonstrate the dependence of LNP physiochemical properties, including size, PDI, shape, and internal structure, on mixing techniques and dialysis steps. We compared eight different mixing approaches with varying FRR (1:1 and 3:1) and laminar/turbulent types. We observed that the 3:1 mixers (Ignite (laminar) and MIVM 3:1) had the smallest sizes (*D*_*h*_) of LNPs but with higher PDI (Sec. 2.1), and a reduction in PDI was observed post-dialysis. The hand-mixed technique had the lowest PDI throughout. In terms of encapsulation efficiency (EE), mixers operating at 3:1 FRR and hand mixing performed better (EE>80%). We observe strong similarity in the performance of the 3:1 MIVM mixers and Ignite. The Ignite-based LNPs also resemble a more spherical shape as compared to those from all other mixing approaches (Sec. 2.2). In terms of internal organization post-dialysis, we observed the hand mixed having the most tightly packed, LC phase (L*α* dominated), followed by MIVM 3:1 (opposite), MIVM 2:2 (adjacent), MIVM 3:1 (adjacent), CIJ, Ignite, MIVM 2:2 (opposite). It also should be noted that dialyzing significantly affected the LC phase packing, shifting from the existence of a L_*α*_/H_*II*_ mixture to a L_*α*_ dominated (See Figs. 2a and b). In conclusion, we observed a strong similarity between both the configurations of the MIVM 3:1 and Ignite post dialysis. Although it should be noted, right after mixing (non-dialyzed LNPs), we did observe some different trends. There were also significant differences observed between the 3:1 and 2:2 MIVMs. This points to strong effects of the FRR on the overall LNP (See Figs. 3a-d). CIJ also had consistent differences compared to the other mixers. This suggests effects arising from the impinging jet mixing mechanism within the 1:1 FRR. Additionally, we concluded from the CFD analysis that the CIJ has two orders of magnitude higher maximum turbulent kinetic energy as compared to MIVM. This also results in poor mixing at the jet impingement for the CIJ. We also reported and concluded in section 2.3 that the fluid dynamics differences affect the mixing timescales, providing some basis for the differences observed in the LNP physicochemical properties. Future study of the mixing process, and pairing with biological performance and scalability studies, is required to better design LNPs with more informed mixer selection at the lab scale. This will ensure LNP formulations with consistent physicochemical properties and easier translation and adoption at larger scales.

## 4 EXPERIMENTAL SECTION

### 4.1 Materials

6-((2-hexyldecanoyl)oxy)-N-(6-((2-hexyldecanoyl)oxy)hexyl)-N-(4-hydroxybutyl)hexan-1-aminium (ALC-0315), 1,2-distearoyl-sn-glycero-3-phosphocholine (DSPC), cholesterol, and 1,2-dimyristoyl-rac-glycero-3-methoxypolyethylene glycol-2000 (DMG-PEG2000) were purchased from Avanti Polar Lipids (Alabama, USA). Purified Torulla Ambion yeast RNA was purchased from Thermo Fisher Scientific (Massachusetts, USA). Glacial acetic acid was purchased from Fisher Chemical (Pennsylvania, USA). Sodium acetate anhydrous, 200 proof ethanol, and RNAse-free water were purchased from Fisher BioReagents (Pennsylvania, USA).

### 4.2 Lipid nanoparticle preparation

An organic solution containing lipids in ethanol was prepared at a total lipid concentration of 6 mg/ml following a 50:38.5:10:1.5 molar ratio of ALC-0315, DSPC, cholesterol, and DMG-PEG_2000_, respectively (See Fig. 1a). Nucleic acid was dissolved in an acetate buffer at pH 5. Five different mixing techniques were utilized, namely, Hand mixing (HM), NanoAssemblr^™^ Ignite (Precision Nanosystems), Nova Impinging Jets Mixer (Helix Biotech), Confined Impinging Jets Mixer (CIJ), and the Multi-Inlet Vortex Mixer (MIVM). The aqueous solution contained 0.1 mg/mL of yRNA for mixers operated at a 3:1 aqueous-to-organics flow rate ratio (i.e., HM, Ignite, and MIVM 3:1), and 0.3 mg/mL of yRNA for mixers operated at equal parts of aqueous and organics (i.e., CIJ, MIVM 2:2, and Helix). The organic and aqueous solutions were then mixed in mixers of interest with the respective flow rate ratios as described in Table 6.

**TABLE 6.** Operating parameters for each mixing technique.

| Mixer | Mixing Type | Flow-Rate Ratio (FRR) | Total-Flow Rate (TFR) (mL/min) |
| --- | --- | --- | --- |
| Hand mixing | Pipette | 3:1 | – |
| Ignite | Toroidal | 3:1 | 12 |
| Helix | Impinging jets | 1:1 | 8 |
| CIJ | Impinging jets | 1:1 | 120 |
| MIVM | Vortex | 1:1 | 160 |
| MIVM | Vortex | 3:1 | 160 |

The nanoparticles were quenched after 5 seconds of exiting the outlet using a 10 mM acetate buffer at pH 5. After quenching, all yRNA-LNPs produced across all mixers had the same lipid concentration of 0.6 mg/mL, yRNA concentration of 30 ug/mL, N/P ratio of 5, lipid/nucleic acid (w/w) of 20, in 10% ethanol. Ethanol in LNP was removed by dialyzing against 10 mM Acetate Buffer at pH 5 using Spectra Por S/P 1 Dialysis Membrane (6-8 kDa MWCO) for 4 hours (Fig. 1b). The antisolvent/buffer pH was kept constant to avoid pH change effects, such as vesicle fusion and size growth. For cryo-preservation purposes, 10% (w/v) of sucrose was added to both the pre- and post-dialyzed LNP formulations, then the samples were shipped to the ANSTO facility.

### 4.3 Dynamic light scattering and zeta potential

Malvern Panalytical Dynamic Light Scattering Zetasizer Pro (Malvern, UK) was utilized for size, polydispersity index (PDI), and zeta potential measurement. The samples were diluted 1:10 to 1 mL using 10 mM acetate buffer, pH 5, and placed into Fisherbrand™ Disposable Cuvettes Semi-micro 1.5 mL for size and polydispersity index measurement. For zeta potential measurement, the samples were diluted 1:10 to 1 mL using 20 mM NaCl to obtain a conductivity of 2 mS/cm, and inserted into Malvern Panalytical Folded Capillary Zeta Cell DTS1070 (Malvern, UK). The method was loaded in the ZS Xplorer software, utilizing water as a dispersant for three runs of 15 measurements taken for each sample and averaged.

### 4.4 Measuring LNP encapsulation

Encapsulation efficiency was measured using the Quant-it™ RiboGreen Assay Kit (Thermo Fisher Scientific, USA) assay. Two sets of controls of the standard curve of the yRNA were prepared in which the amount of RiboGreen dye was fixed at 0. 25% v / v and both had the standard yRNA (4 µg/mL) serially diluted to produce the final yRNA of 0, 0.1, 0.2, 0.4, 0.6, 0.8, and 1 µg/mL, with the balance of the volume adjusted using 1× TE buffer. The first series, designated as the control without Triton X-100 (Fisher Scientific, USA), contained only yRNA, TE buffer, and RiboGreen dye. The second series, designated as control with Triton X-100, was prepared in the same manner except that a constant amount of 0.25% Triton X-100 was included in each sample, replacing an equivalent volume of TE buffer to maintain equal total volume.

LNP formulations were diluted to 8 µg/mL (based on yRNA content) in 1× TE buffer were assayed in parallel with and without Triton X-100. The first series, designated as the sample control without Triton X-100, contained 7.5% LNP sample, 42.5% TE buffer, and 0.25% v/v RiboGreen dye. The second series, designated as sample control with Triton X-100, was prepared using the same amount of LNP sample but with 0.25% Triton X-100 solution added; the volume of the TE buffer was reduced to 17. 5% to maintain the same total volume.

All samples, including standards and LNPs, were gently vortexed, and 100 µL of each was transferred to wells of a black, flat bottom, half-area, nontreated 96-well microplate (CorningTM, USA). The calibration curve samples were transferred in duplicates, while the LNP samples were transferred in triplicate. The plates were incubated in a dark environment at room temperature for 5 minutes prior to measurement. Fluorescence was recorded at an excitation wavelength of 485 nm and an emission wavelength of 528 nm using a SpectraMax i3x multimode microplate reader (Molecular Devices, San Jose, CA, USA). Two replicates were conducted for calibration samples, and three replicates were conducted for each LNP sample.

### 4.5 Synchrotron SAXS measurements

SAXS measurements were primarily performed at the Small and Wide Angle X-ray Scattering (SAXS/WAXS) beamline of the Australian Synchrotron, part of ANSTO ^64^. The autoloader sample environment developed at the Australian Synchrotron was used at the room temperature (300 K) of the SAXS/WAXS experimental hutch. Scattering at low q values, *q* = 4*π* sin *θ*/*λ*, where *λ* is the X-ray wavelength is 1.0332 Å and 2θ is the scattering angle) were recorded at a sample–detector distance (SDD) of 5.0 m with a photon energy of 7 keV, producing a q range of 0.0028 < *q* < 3.46 Å^−1^. Cryo-preserved LNPs at –80^°^C were thawed at 4^°^C and background buffers. LNP samples were then concentrated using Amicon Ultra centrifugal filters (50 kDa MWCO, Milipore Sigma) on a centrifuge operated at 500 rcf. The filters were initially primed with 30% ethanol once, followed by two rinses with RNAse-free water, each at 500 rcf for 2 minutes. The final lipid concentration in each sample was around 3 mg/mL. Concentrated LNPs and background buffers (100 uL aliquots) were loaded into 96-well plates. The samples were then remotely drawn sequentially into a quartz capillary held stationary in the beam, and up to 15 scattering measurements were performed on each sample as the sample flowed through the capillary and was ejected back into the sample well. The capillary was washed with water and 2.0% Hellmanex detergent solution between samples. 2d scattering patterns were radially integrated into 1D scattering functions, i.e., *I*(*q*), using the in-house developed software package ScatterBrain (ver 2.82). The resulting SAXS profile follows, *I*(*q*) = ν(Δ*ρ*)*P*(*q*)*S*(*q*). Here, ν, Δ*ρ*, and *P*(*q*) are the volume fraction, difference in the scattering length density (SLD) between LNPs and the buffer, and the intra-particle form factor of the LNPs, respectively. In our experiments, we assume *S*(*q*) = 1, as there is no inter-particle structure factor. The scattering intensities were plotted on an absolute scale with units of *cm*^−1^, with water as a calibration standard.

### 4.6 SAXS data processing and analysis

The reduced data was loaded into PRIMUS from ATSAS 4.0.11^65^. Each of the 15 data sets for each LNP and the background buffers was averaged. Finally, the LNP SAXS profiles were subtracted for their respective background buffers, making the SAXS profile follow, *I*(*q*)_*LNP*_ = *I*(*q*) – *I*_*buffer*_ = ν(Δ*ρ*)*P*(*q*)_*LNP*_*S*(*q*)_*LNP*_. The background-subtracted 1d-SAXS profiles were then loaded onto the SASView 5.0.6 fitting interface. For the Porod exponent (*p*), *I*(*q*) was fitted using a power law for 0.01 < q < 0.1 Å^−1^. Shape-independent Broad-Lorentzian type peak was utilized to extract the peak features for both the non-dialyzed and dialyzed formulations from each mixing approach ^66^. This is described by the following equation,

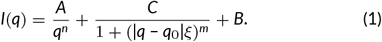

Here, the peak location is related to the d-spacing as *q*_0_ = 2*π*/*d*_0_ (Bragg law), *A* is the Porod law scale factor, *n* the Porod exponent, *C* is the Lorentzian scale factor, *m* the exponent of *q, ξ* the screening length, and *B* the flat background. It should be noted, the Porod exponent, *n*, here is for the BL model and should not be confused with *p* from the power law fit within the q range of 0.01 to 0.1 Å^−1^. The peak location, i.e., *q*_0_, was given a fixed range around the LC phase peaks, and an initial fit was performed with the Levenberg–Marquardt (LM) algorithm. For the *m* and *n* values, the ranges were limited to 0 to 6 and 0 to 4. Finally, the Differential Evolution Adaptive Metropolis (DREAM) algorithm was run on the initialized LM fit results.

The inversion for extracting the Pairwise Distance Distribution Function (PDDF) was performed in GNOM GUI available through the BioXTAS RAW 2.3.0^67^. PDDF was preferred over Guinier fitting to extract *Rg*, because of the non-sphericity and limited low q data points ^39^. Each 1d-SAXS profiles were loaded in the Indirect Fourier Transform (IFT), and PDDF was calculated using GNOM. Each iteration was chosen with a manual guess of the *Dmax* until a reasonably good solution (Total estimate) was reached with a low *χ*^2^. The PDDF constraint of being zero at *Dmax* values was switched off.

For the 3d-reconstructions of the electron density maps, the DENSS algorithm was utilized ^59^. The calculated scattering intensity is

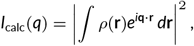

and the electron density is refine d by minimizi ng,

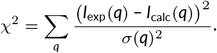

DENSS was run in slow mode using the largest axis as the symmetry, with the enantiomer selection turned on. Reconstructions were visualized using four contour density levels, which were represented by *σ*, i.e., the standard deviations above the mean electron density value generated in the model. The color contours used were: 15σ (red), 10σ (green), 5σ (cyan), and 2σ (blue).

Volatility of Ratio (*Vr* ) metric was calculated as follows,

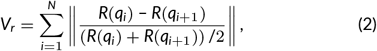

where *R* is the ratio of intensities at *q*_*i*_ ^62^. This was applied to the *I*(*q*) profiles altogether. From these heat maps of Vr were generated in a clustered form, where more similar cases would be closer to each other. The heat maps for the whole q range and q range fixed from 0.08 to 0.2 Å^−1^ were calculated to compare the similarity of the LC phase separately. For the force plot visualizations, the size of the nodes/circles is proportional to the *Rg* of the SAXS profile. The distance between two nodes/circles is directly proportional to the Vr similarity of the two.

### 4.7 Computational fluid dynamics simulations

For the CFD simulations, we used an in-house solver named Multiphysics Finite Element Solver (MUPFES) ^68–70^. The solver uses the residual-based variational multiscale finite element method to solve the Navier-Stokes equation, and a streamline-upwind stabilized finite element method to solve the advection-diffusion equation. The solver has been extensively validated and employed to simulate complex turbulent flows in various peer-reviewed papers ^71,72^. The exact numerical details and simulation setup can be found in Jia *et al*. ^63^

## AUTHOR CONTRIBUTIONS

H.M., T.B., S.R.D., K.D.R., and A.M.A. designed all experiments. T.B. formulated all LNPs and performed the DLS experiments and EE quantifications. H.M., T.B., P.M.S., L.S.M., and I.M. performed the synchrotron SAXS experiments. H.M. and P.M.S. performed the SAXS data curation, processing, and analysis. Simulations were performed and postprocessed by D.J. and M.M. H.M., T.B., D.J., M.M., K.D.R., and A.M.A. wrote the original draft. K.D.R. and A.M.A reviewed and edited the manuscript. K.D.R. and A.M.A. were responsible for project administration, funding acquisition, and supervision.

## ACKNOWLEDGMENTS

This research was supported by a grant from Eli Lilly and Company (USA), part of the Eli Lilly and Purdue University Research Alliance Center (LPRC). The SAXS experiments for this work were conducted on the SAXS/WAXS beamline of the Australian Synchrotron in Melbourne, part of ANSTO (Award reference #: AS243/SAXS/22446, awarded to K.D.R.). We thank Faezeh Masoomi, Gabriel Harris, Mojhdeh Baghban-bashi, Peyman Dastyar, and Sophia R. Dasaro in assistance with the LNP preparations. The authors would like to thank Anas Aljabbari for insightful discussions. This work benefited from the use of the SasView application, originally developed under NSF award DMR-0520547. SasView contains code developed with funding from the European Union’s Horizon 2020 research and innovation programme under the SINE2020 project, grant agreement No. 654000.

## 5 DATA AVAILABILITY STATEMENT

The data supporting this study’s findings are available from the corresponding author upon reasonable request.

## SUPPORTING INFORMATION

Supporting Information is available from the Wiley Online Library or from the author.

## REFERENCES

1. Cullis PR, Hope MJ. Lipid nanoparticle systems for enabling gene therapies. Molecular Therapy. 2017;25(7):1467–1475.

2. Hou X, Zaks T, Langer R, Dong Y. Lipid nanoparticles for mRNA delivery. Nature Reviews Materials. 2021;6(12):1078–1094.

3. Schoenmaker L, Witzigmann D, Kulkarni JA, Verbeke R, Kersten G, Jiskoot W, et al. mRNA-lipid nanoparticle COVID-19 vaccines: Structure and stability. International journal of pharmaceutics. 2021;601:120586.

4. Mitchell MJ, Billingsley MM, Haley RM, Wechsler ME, Peppas NA, Langer R. Engineering precision nanoparticles for drug delivery. Nature reviews drug discovery. 2021;20(2):101–124.

5. Bian X, Luo Z, Peng B, Chen J, Lo PK, Zhou L, et al. Engineered Bionanomaterials for Precision Delivery of Nucleic Acid Drugs. Small. 2025;p. e02667.

6. Zong Y, Lin Y, Wei T, Cheng Q. Lipid nanoparticle (LNP) enables mRNA delivery for cancer therapy. Advanced Materials. 2023;35(51):2303261.

7. Eygeris Y, Patel S, Jozic A, Sahay G. Deconvoluting lipid nanoparticle structure for messenger RNA delivery. Nano letters. 2020;20(6):4543–4549.

8. Yanez Arteta M, Kjellman T, Bartesaghi S, Wallin S, Wu X, Kvist AJ, et al. Successful reprogramming of cellular protein production through mRNA delivered by functionalized lipid nanoparticles. Proceedings of the National Academy of Sciences. 2018;115(15):E3351–E3360.

9. Ripoll M, Martin E, Enot M, Robbe O, Rapisarda C, Nicolai MC, et al. Optimal self-assembly of lipid nanoparticles (LNP) in a ring micromixer. Scientific Reports. 2022;12(1):9483.

10. Maeki M, Fujishima Y, Sato Y, Yasui T, Kaji N, Ishida A, et al. Understanding the formation mechanism of lipid nanoparticles in microfluidic devices with chaotic micromixers. PLoS One. 2017;12(11):e0187962.

11. Shepherd SJ, Warzecha CC, Yadavali S, El-Mayta R, Alameh MG, Wang L, et al. Scalable mRNA and siRNA lipid nanoparticle production using a parallelized microfluidic device. Nano letters. 2021;21(13):5671–5680.

12. Sealy A. Manufacturing moonshot: How Pfizer makes its millions of Covid-19 vaccine doses. CNN: CNN Health. 2021;.

13. Dasaro SR, Singh A, Vlachos P, Ristroph KD. Mechanistic insights into how mixing factors govern polyelectrolyte-surfactant complexation in RNA lipid nanoparticle formulation. Journal of Colloid and Interface Science. 2025;678:98–107.

14. Evers MJ, Kulkarni JA, van der Meel R, Cullis PR, Vader P, Schiffelers RM. State-of-the-art design and rapid-mixing production techniques of lipid nanoparticles for nucleic acid delivery. Small methods. 2018;2(9):1700375.

15. Pratsinis A, Fan Y, Portmann M, Hammel M, Kou P, Sarode A, et al. Impact of non-ionizable lipids and phase mixing methods on structural properties of lipid nanoparticle formulations. International Journal of Pharmaceutics. 2023;637:122874.

16. Petersen DMS, Chaudhary N, Arral ML, Weiss RM, Whitehead KA. The mixing method used to formulate lipid nanoparticles affects mRNA delivery efficacy and organ tropism. European Journal of Pharmaceutics and Biopharmaceutics. 2023;192:126– 135.

17. Jürgens DC, Deßloch L, Porras-Gonzalez D, Winkeljann J, Zielinski S, Munschauer M, et al. Lab-scale siRNA and mRNA LNP manufacturing by various microfluidic mixing techniques–an evaluation of particle properties and efficiency. OpenNano. 2023;12:100161.

18. Riewe J, Erfle P, Melzig S, Kwade A, Dietzel A, Bunjes H. Antisolvent precipitation of lipid nanoparticles in microfluidic systems– A comparative study. International journal of pharmaceutics. 2020;579:119167.

19. Mo Y, Keszei AF, Kothari S, Liu H, Pan A, Kim P, et al. Lipid-siRNA Organization Modulates the Intracellular Dynamics of Lipid Nanoparticles. Journal of the American Chemical Society. 2025;147(12):10430–10445.

20. O’Brien Laramy MN, Costa AP, Cebrero YM, Joseph J, Sarode A, Zang N, et al. Process robustness in lipid nanoparticle production: a comparison of microfluidic and turbulent jet mixing. Molecular pharmaceutics. 2023;20(8):4285–4296.

21. Chen X, Li M, Jiang F, Hong L, Liu Z. Revealing a Correlation between Structure and in vitro activity of mRNA Lipid Nanoparticles. bioRxiv. 2024;p. 2024–09.

22. Na GS, Joo JU, Lee JY, Yun Y, Kaang BK, Yang JS, et al. Full-cycle study on developing a novel structured micromixer and evaluating the nanoparticle products as mRNA delivery carriers. Journal of Controlled Release. 2024;373:161–171.

23. Hourdel L, Lebaz N, Peral F, Ripoll M, Briançon S, Bensaid F, et al. Overview on LNP-mRNA encapsulation unit operation: Mixing technologies, scalability, and influence of formulation & process parameters on physico-chemical characteristics. International Journal of Pharmaceutics. 2025;p. 125297.

24. Ristroph KD. Drugs need to be formulated with scale-up in mind. Journal of Controlled Release. 2024;373:962–966.

25. Udepurkar A, Devos C, Sagmeister P, Destro F, Inguva P, Ahmadi S, et al. Structure and Morphology of Lipid Nanoparticles for Nucleic Acid Drug Delivery: A Review. ACS nano. 2025;.

26. Padilla MS, Shepherd SJ, Hanna AR, Kurnik M, Zhang X, Chen M, et al. Solution biophysics identifies lipid nanoparticle nonsphericity, polydispersity, and dependence on internal ordering for efficacious mRNA delivery. bioRxiv. 2025;p. 2024–12.

27. Hassett KJ, Higgins J, Woods A, Levy B, Xia Y, Hsiao CJ, et al. Impact of lipid nanoparticle size on mRNA vaccine immunogenicity. Journal of Controlled Release. 2021;335:237–246.

28. Zhang L, More KR, Ojha A, Jackson CB, Quinlan BD, Li H, et al. Effect of mRNA-LNP components of two globally-marketed COVID-19 vaccines on efficacy and stability. npj Vaccines. 2023;8(1):156.

29. Lam K, Schreiner P, Leung A, Stainton P, Reid S, Yaworski E, et al. Optimizing lipid nanoparticles for delivery in primates. Advanced Materials. 2023;35(26):2211420.

30. Cárdenas M, Campbell RA, Arteta MY, Lawrence MJ, Sebastiani F. Review of structural design guiding the development of lipid nanoparticles for nucleic acid delivery. Current Opinion in Colloid & Interface Science. 2023;66:101705.

31. Hammel M, Fan Y, Sarode A, Byrnes AE, Zang N, Kou P, et al. Correlating the structure and gene silencing activity of oligonucleotide-loaded lipid nanoparticles using small-angle X-ray scattering. ACS nano. 2023;17(12):11454–11465.

32. Bi D, Wilhelmy C, Unthan D, Keil IS, Zhao B, Kolb B, et al. On the Influence of Fabrication Methods and Materials for mRNA-LNP Production: From Size and Morphology to Internal Structure and mRNA Delivery Performance In Vitro and In Vivo. Advanced healthcare materials. 2024;13(26):2401252.

33. Carrasco MJ, Alishetty S, Alameh MG, Said H, Wright L, Paige M, et al. Ionization and structural properties of mRNA lipid nanoparticles influence expression in intramuscular and intravascular administration. Communications biology. 2021;4(1):956.

34. Garaizar A, Díaz-Oviedo D, Zablowsky N, Rissanen S, Köbberling J, Sun J, et al. Toward understanding lipid reorganization in RNA lipid nanoparticles in acidic environments. Proceedings of the National Academy of Sciences. 2024;121(45):e2404555121.

35. Pattipeiluhu R, Zeng Y, Hendrix MM, Voets IK, Kros A, Sharp TH. Liquid crystalline inverted lipid phases encapsulating siRNA enhance lipid nanoparticle mediated transfection. Nature Communications. 2024;15(1):1303.

36. Nong J, Gong X, Dang QM, Tiwari S, Patel M, Wu J, et al. Multistage-mixing to control the supramolecular structure of lipid nanoparticles, thereby creating a core-then-shell arrangement that improves performance by orders of magnitude. bioRxiv. 2025;p. 2024–11.

37. Chen X, Ye Y, Li M, Zuo T, Xie Z, Ke Y, et al. Structural characterization of mRNA lipid nanoparticles (LNPs) in the presence of mRNA-free LNPs. Journal of Controlled Release. 2025;p. 114082.

38. Li Z, Carter J, Santos L, Webster C, Van Der Walle CF, Li P, et al. Acidification-induced structure evolution of lipid nanoparticles correlates with their in vitro gene transfections. ACS nano. 2023;17(2):979–990.

39. Caselli L, Conti L, De Santis I, Berti D. Small-angle X-ray and neutron scattering applied to lipid-based nanoparticles: Recent advancements across different length scales. Advances in Colloid and Interface Science. 2024;327:103156.

40. Cheng MHY, Leung J, Zhang Y, Strong C, Basha G, Momeni A, et al. Induction of bleb structures in lipid nanoparticle formulations of mRNA leads to improved transfection potency. Advanced Materials. 2023;35(31):2303370.

41. Kulkarni JA, Darjuan MM, Mercer JE, Chen S, Van Der Meel R, Thewalt JL, et al. On the formation and morphology of lipid nanoparticles containing ionizable cationic lipids and siRNA. ACS nano. 2018;12(5):4787–4795.

42. Roces CB, Lou G, Jain N, Abraham S, Thomas A, Halbert GW, et al. Manufacturing considerations for the development of lipid nanoparticles using microfluidics. Pharmaceutics. 2020;12(11):1095.

43. Johnson BK, Prud’homme RK. Chemical processing and micromixing in confined impinging jets. AIChE Journal. 2003;49(9):2264–2282.

44. Markwalter CE, Prud’homme RK. Design of a small-scale multi-inlet vortex mixer for scalable nanoparticle production and application to the encapsulation of biologics by inverse flash nanoprecipitation. Journal of pharmaceutical sciences. 2018;107(9):2465–2471.

45. Liu Y, Cheng C, Prud’homme RK, Fox RO. Mixing in a multi-inlet vortex mixer (MIVM) for flash nano-precipitation. Chemical Engineering Science. 2008;63(11):2829–2842.

46. Devos C, Mukherjee S, Inguva P, Singh S, Wei Y, Mondal S, et al. Impinging jet mixers: A review of their mixing characteristics, performance considerations, and applications. AIChE Journal. 2025;71(1):e18595.

47. Hardianto A, Muscifa ZS, Widayat W, Yusuf M, Subroto T. The effect of ethanol on lipid nanoparticle stabilization from a molecular dynamics simulation perspective. Molecules. 2023;28(12):4836.

48. Maeki M, Kimura N, Okada Y, Shimizu K, Shibata K, Miyazaki Y, et al. Understanding the effects of ethanol on the liposome bilayer structure using microfluidic-based time-resolved small-angle X-ray scattering and molecular dynamics simulations. Nanoscale Advances. 2024;6(8):2166–2176.

49. Zhao L, Xu Z, Li H, Liu L, Chen S, Peng Z, et al. A review of confined impinging jet reactor (CIJR) with a perspective of mRNA-LNP vaccine production. Reviews in Chemical Engineering. 2024;40(8):887–916.

50. Peng H, Li Z, Cai Z, Gao Z. Study of a Novel Method to Weaken the Backmixing in a Multi-Inlet Vortex Mixer. Processes. 2024;12(3):476.

51. Vargas R, Romero M, Berasategui T, Narváez-Narváez DA, Ramirez P, Nardi-Ricart A, et al. Dialysis is a key factor modulating interactions between critical process parameters during the microfluidic preparation of lipid nanoparticles. Colloid and Interface Science Communications. 2023;54:100709.

52. Terada T, Kulkarni JA, Huynh A, Chen S, van der Meel R, Tam YYC, et al. Characterization of lipid nanoparticles containing ionizable cationic lipids using design-of-experiments approach. Langmuir. 2021;37(3):1120–1128.

53. Gilbert J, Sebastiani F, Arteta MY, Terry A, Fornell A, Russell R, et al. Evolution of the structure of lipid nanoparticles for nucleic acid delivery: From in situ studies of formulation to colloidal stability. Journal of colloid and interface science. 2024;660:66–76.

54. Johansen D, Trewhella J, Goldenberg DP. Fractal dimension of an intrinsically disordered protein: Small-angle X-ray scattering and computational study of the bacteriophage λ N protein. Protein Science. 2011;20(12):1955–1970.

55. Schmidt PW. Small-angle scattering studies of disordered, porous and fractal systems. Applied Crystallography. 1991;24(5):414–435.

56. Shah RM, Mata JP, Bryant G, de Campo L, Ife A, Karpe AV, et al. Structure analysis of solid lipid nanoparticles for drug delivery: a combined USANS/SANS study. Particle & Particle Systems Characterization. 2019;36(1):1800359.

57. Beaucage G, Kammler HK, Pratsinis SE. Particle size distributions from small-angle scattering using global scattering functions. Applied Crystallography. 2004;37(4):523–535.

58. Kulkarni JA, Witzigmann D, Chen S, Cullis PR, Van Der Meel R. Lipid nanoparticle technology for clinical translation of siRNA therapeutics. Accounts of chemical research. 2019;52(9):2435– 2444.

59. Grant TD. Ab initio electron density determination directly from solution scattering data. Nature methods. 2018;15(3):191–193.

60. Dao HM, AboulFotouh K, Hussain AF, Marras AE, Johnston KP, Cui Z, et al. Characterization of mRNA lipid nanoparticles by electron density mapping reconstruction: x-ray scattering with density from solution scattering (DENSS) algorithm. Pharmaceutical Research. 2024;41(3):501–512.

61. Graewert MA, Wilhelmy C, Bacic T, Schumacher J, Blanchet C, Meier F, et al. Quantitative size-resolved characterization of mRNA nanoparticles by in-line coupling of asymmetrical-flow field-flow fractionation with small angle X-ray scattering. Scientific reports. 2023;13(1):15764.

62. Hura GL, Budworth H, Dyer KN, Rambo RP, Hammel M, McMurray CT, et al. Comprehensive macromolecular conformations mapped by quantitative SAXS analyses. Nature methods. 2013;10(6):453–454.

63. Jia D, Majidi M, Ristroph KD, Ardekani A. High-Fidelity Simulations of Two Miscible Fluids in Small Scale Turbulent Mixers Using a Variational Multiscale Finite Element Method. arXiv e-prints. 2025;p. 2509.12029.

64. Kirby NM, Mudie ST, Hawley AM, Cookson DJ, Mertens HD, Cowieson N, et al. A low-background-intensity focusing smallangle X-ray scattering undulator beamline. Applied Crystallography. 2013;46(6):1670–1680.

65. Manalastas-Cantos K, Konarev PV, Hajizadeh NR, Kikhney AG, Petoukhov MV, Molodenskiy DS, et al. ATSAS 3.0: expanded functionality and new tools for small-angle scattering data analysis. Applied Crystallography. 2021;54(1):343–355.

66. The SasView Team. SasView Version 6.0.1 Documentation 2019. Available from: https://www.sasview.org/docs/user/qtgui/Perspectives/Fitting/plugin.html.

67. Hopkins JB. BioXTAS RAW 2: new developments for a free open-source program for small-angle scattering data reduction and analysis. Applied Crystallography. 2024;57(1):194–208.

68. Moghadam ME, Vignon-Clementel IE, Figliola R, Marsden AL, of Congenital Hearts Alliance (MOCHA) Investigators M, et al. A modular numerical method for implicit 0D/3D coupling in cardiovascular finite element simulations. Journal of Computational Physics. 2013;244:63–79.

69. Esmaily-Moghadam M, Bazilevs Y, Marsden AL. A bipartitioned iterative algorithm for solving linear systems arising from incompressible flow problems. Computer Methods in Applied Mechanics and Engineering. 2015;286:40–62.

70. Esmaily-Moghadam M, Bazilevs Y, Marsden AL. A new preconditioning technique for implicitly coupled multidomain simulations with applications to hemodynamics. Computational Mechanics. 2013;52(5):1141–1152.

71. Jia D, Esmaily M. Characterization of the ejector pump performance for the assisted bidirectional Glenn procedure. Fluids. 2022;7(1):31.

72. Jia D, Lee Baker J, Rameau A, Esmaily M. Simulation of a vacuum helmet to contain pathogen-bearing droplets in dental and otolaryngologic outpatient interventions. Physics of Fluids. 2021;33(1).

